# Predicting temozolomide response in low-grade glioma patients with large-scale machine learning

**DOI:** 10.1101/2024.12.22.629995

**Authors:** Hanqin Du, Chayanit Piyawajanusorn, Ghita Ghislat, Pedro J. Ballester

**Affiliations:** Department of Bioengineering, Imperial College, London, United Kingdom

**Keywords:** Temozolomide, low-grade glioma, machine learning, drug response prediction, tumor omics

## Abstract

**Background:** Temozolomide is the primary chemotherapeutic agent and first-line treatment for low-grade glioma (LGG). Although LGGs are generally less aggressive than high-grade gliomas (HGGs), they can eventually progress into HGGs, making it crucial to maximise the efficacy of initial LGG treatment.

**Methods:** In this study, we analysed data from 109 LGG patients in The Cancer Genome Atlas (TCGA) to assess the predictive performance of 12 machine learning (ML) classification algorithms in forecasting temozolomide response, utilising six types of omics data. Cross-validation and bootstrapping bias correction were employed to compare the performance of these models with that of a conventional biomarker-based model using MGMT promoter methylation status.

**Results:** Among the models, the miRNA-based approach using the XGBoost algorithm showed the most promising predictive performance, with a Matthews Correlation Coefficient (MCC) of 0.447, outperforming the auto-ML method JADBio (MCC = 0.250) and the MGMT biomarker-based model. Converting the task into a regression framework by encoding cancer response as a continuous variable generally weakened predictive performance, with best model based on methylation profile (MCC = 0.344). Additionally, ML models focusing on biomarker-relevant features demonstrated improved prediction accuracy, though not to the extent of omics-based models, with the support vector machine (SVM) model achieving an MCC of 0.331. Incorporating clinical variables, such as patient age and Karnofsky score, further enhanced predictive power, with the LR-OMC model achieving the highest MCC (0.483). Feature importance analysis identified six significant miRNA factors, including three tumour-related miRNAs (miR-335, let-7f, and miR-7-2) and three potential biomarkers (miR-204, miR-6513, and miR-376).

**Discussion:** Overall, this study systematically demonstrates the potential of large-scale analyses combining machine learning and omics data to predict temozolomide response, offering superior predictive accuracy compared to standard MGMT biomarkers.

## INTRODUCTION

Glioma is a type of brain tumour originating in glial cells, which surround neurons and facilitate their function^1^. In adults, gliomas account for 80% of primary malignant tumours in the central nervous system^2^. Based on histopathological features and proliferation rates, gliomas are graded into low-grade gliomas (LGG) and high-grade gliomas (HGG). LGG accounts for approximately 30% of gliomas^3^, with astrocytoma, oligodendroglioma and ganglioglioma being the most common subtypes. Compared to HGGs, LGG cells exhibit a higher resemblance to normal cells and proliferate more slowly, which makes it more difficult to evaluate their growth. Nevertheless, finding an effective drug treatment is remains critical as LGGs may develop into HGGs.

Surgical resection has been establishing as the mainstay of first-line treatment for LGG patients, as several studies have shown that a greater extent of resection correlates with a better prognosis. However, the infiltrative nature of LGGs often poses a challenge to surgery, as these tumours may extend into critical brain regions that are difficult to access, thereby hindering gross total resection challenging^4,5^. Radiotherapy and chemotherapy, administered as monotherapy or in combination, are common adjuvant options to surgery, despite being associated with side effects and limitations, such as toxicity and resistance.

Temozolomide is a first-line chemotherapeutic agent for various brain cancer types, including LGG. As a prodrug, temozolomide is chemically unstable at physiological pH (above pH7), decomposing into monomethyl triazene 5-(3-methyltriazen-1-yl)-imidazole-4-carboxamide (MTIC)^6^. MTIC further hydrolyses to 5-aminoimidazole-4-carboxamide (AIC) and a methyldiazonium cation, the latter of which methylates DNA at N7 positions of guanine, leading to DNA replication inhibition and cell death. Given that brain tumours often exhibit a slightly alkaline microenvironment compared to surrounding healthy tissue, temozolomide preferentially targets tumour cells rather than normal cells^7^. While temozolomide has been proven effective in extending the lifespan of patients with gliomas, quality of life remains a significant concern. Various toxicities associated with temozolomide has been reported, such as nausea, vomiting, anorexia, and hematologic toxicities. Due to the heterogeneity of cancer cells, the effectiveness of temozolomide varies among patients. Thus, accurately predicting drug response before treatment is crucial to minimize unnecessary side effects.

The methylation status of the O6-methylguanine-DNA methyltransferase (MGMT) promoter is the most widely recognized biomarker for predicting temozolomide resistance^8^. MGMT plays a crucial role in temozolomide resistance by directly reversing the damage induced by temozolomide^9^. Although assessing MGMT methylation is generally recommended in clinical practice, its utility is limited due to a significant incidence of false positives^10^, and the MGMT methylation status may change over the course of treatment^11,12^. In addition to MGMT, multiple DNA repair pathways can influence the response to temozolomide. For instance, when MGMT fails to repair the DNA damage, the DNA mismatch repair (MMR) system attempts to repair temozolomide-induced O6-meG the temozolomide-induced mismatches by recognizing the damage and excising the newly synthesized DNA strand. This leads to repetitive repair cycles which ultimately cause cell cycle arrest and apoptosis. When the MMR pathway is damaged or down-regulated, DNA synthesis and cell cycle will continue, leading to resistance to temozolomide^13^. Therefore, defects in the MMR system and/or the expression of MGMT can also contribute to a tumor’s resistance to temozolomide. Previous research by Dunn et al. and Karayan-Tapon et al. demonstrated that elevated MGMT expression or decreased methylation of the MGMT promoter, which leads to increased MGMT levels, is associated with reduced overall survival in glioma patients undergoing temozolomide treatment^8,14^. In addition, base excision repair (BER) and nucleotide excision repair (NER) pathways may also repair temozolomide-induced DNA damage, while the process is more complex and less efficient than the MGMT repair mechanism and requires the participation of multiple proteins^15-17^. Beyond DNA repair mechanism, the activation of anti-apoptotic pathway, downregulation of tumor suppressor proteins, and other mechanisms can indirectly enhance cancer cell drug resistance^18,19^. Therefore, examining multiple omics profiles per sample rather than restricting to a univariate MGMT marker is expected to lead to higher prediction accuracy.

Machine learning (ML) has been shown to be effective in leveraging matched drug sensitivity and omics data to predict the response of unseen cell lines to a given drug^20-24^. Such drug sensitivity is summarised by the half-maximal inhibitory concentration (IC_50_) of the drug acting on the cell line. Several studies employed supervised learning algorithms such as Support Vector Machine (SVM), Random Forest (RF), and Neural network (NN) using omics data including mRNA expression, copy number variation (CNV), DNA methylation, and gene mutations to predict cancer cell sensitivity to a given drug^22,25-27^. Although cell lines have been extensively used to study anticancer agents and to establish vast molecular and drug response datasets, it remains unclear how well they represent the biological characteristics and drug responses of patient tumor samples e.g., due to the lack of a microenvironment in these models. Moreover, many studies have pointed out that there are differences between cell-line and tissue samples in terms of gene expression, protein expression, CNV, and other omics. More importantly, the response of a cell line to a drug can be extremely different from the response of a patient to the same drug^28^. Thus, even when drug resistance is accurately predicted in cell lines, cell-line drug response predictors are not well-represented when applied to predict the sensitivity of tumor patients to certain drugs, demonstrating that their performance often declines^29-35^.

The growing availability of matched drug-response and molecular-profiled tumour data of patients provides an unprecedented opportunity to perform ML analysis in precision medicine. The Cancer Genome Atlas (TCGA), accessed through the Genomic Data Commons (GDC) database, comprising multiple molecular profiles and clinical datasets for more than 11,000 patients across over 33 cancer types^36,37^. This project provides many cases with detailed clinical records, and follow-up records, as well as a variety of molecular profiles, including mRNA expression level, miRNA and isomiR expression level, methylation beta value and copy number variation (CNV)^38^. Although clinical pharmaco-omic data are more relevant than preclinical data, they are significantly scarcer. This scarcity poses a challenge for data-hungry deep learning models, which typically require large datasets to achieve optimal performance. Consequently, the application of deep learning approaches may be less promising in this context^39^. In recent years, many studies have been devoted to predicting patients’ responses to drugs using various tumor molecular profiles^40-43^. To the best of our knowledge, no prior study has applied a comprehensive range of ML algorithms to classify LGG patients’ responses to temozolomide using diverse molecular profiles and clinical datasets.

To investigate this question, we retrieved data for six molecular profiles and clinical information for each of the 109 temozolomide-treated patients with valid records in the TCGA project. The molecular profiles included miRNA, isomiR, mRNA expression level measured in FPKM and FPKM-UQ, DNA methylation, and copy number variation. We evaluated six widely used ML algorithms, including both classification and regression models, with and without Optimal Model Complexity (OMC) feature selection strategy. This approach produced a total of 12 models evaluated on each type of the six molecular profiles to predict LGG patients’ response to temozolomide. The goal of this study was to identify robust ML models capable of classifying LGG patients into temozolomide responders or non-responders. The predictive performances of these models were compared that of the single-gene marker MGMT methylation status and an auto-ML method. Overall, we found that employing appropriate data and a range of algorithms enhances model accuracy, provides direction for exploring advanced ML algorithms and has the potential to guide clinicians in identifying glioma patients’ responses to maximize treatment efficacy while minimizing side effects.

## RESULTS

### Experimental design for the systematic ML analysis

We obtained records of 134 LGG patients with comprehensive omics data from the TCGA project through the GDC protocol. After filtering, we retained records for 109 of these patients with a history of temozolomide administration and valid drug response (Figure S1). Our study encompassed six omics data types: mRNA expression in fragments per kilobase of transcript per million fragments mapped (FPKM), mRNA upper-quartile expression (FPKM-UQ), miRNA and isomiR expression in reads per million (RPM), Copy number variation and methylation beta value. After constructing a dataset from each omics profile, we evaluate each of them with 12 ML algorithms. For each algorithm-omics pair, we used five different random seeds to carry out five nested Cross-Validation (CV) runs. Model performance was compared based on the median of Matthews Correlation Coefficient (mMCC), Area Under the Curve of Precision-Recall (mPR-AUC) and Receiver Operating Characteristic curve (mROC-AUC) (Figure 1).

**Figure 1.**
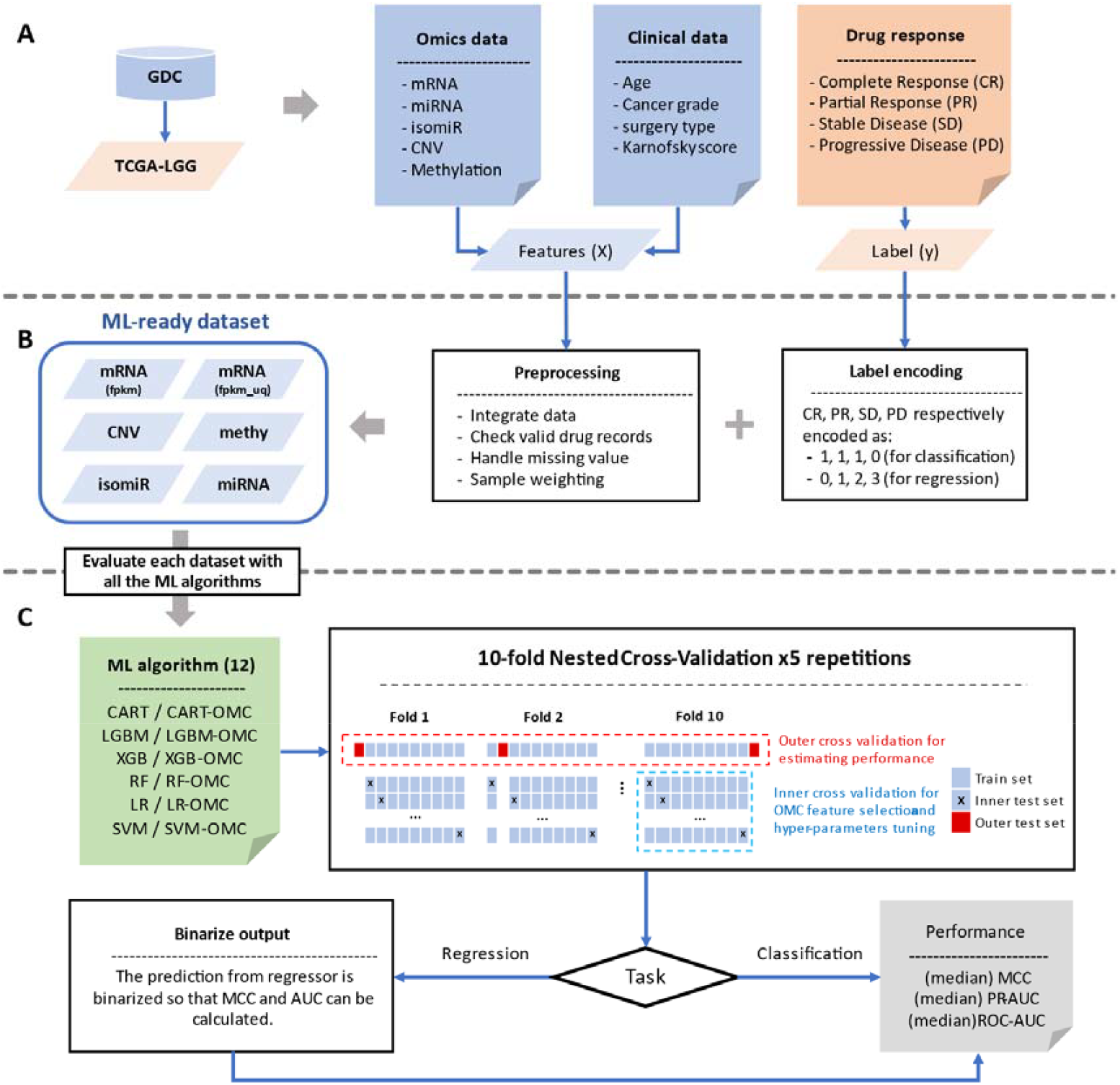
Overview of the systematic ML analysis. **A** Cancer patients’ data were obtained from TCGA and the omics data and drug response were extracted. **B** The omics data and clinical records are used as features. These features were first filtered and pre-processed, then integrated with encoded drug responses to build 6 datasets, where each corresponding to a specific type of omics data. The clinical records are considered optional features that have the potential to enhance the overall performance of the model. **C** The prediction performance of each dataset was evaluated using 12 machine learning algorithms and evaluated using 5 NCV runs (each repeated with a different random seed). For the regression task, the threshold that maximizes the performance on the train set is used for binarized continuous prediction results to calculate MCC and AUC for comparison.

All 109 patients but one had undergone prior surgical intervention. Specifically, 55 patients underwent subtotal resection (SR), while 53 underwent gross total resection (GTR). The sample for the patient without any surgical resection was obtained with a biopsy. According to the National Cancer Institute (NCI) Thesaurus (https://docs.gdc.cancer.gov/Data_Dictionary/gdcmvs/), the terms SR and GTR correspond to the NCI Thesaurus Code C131680 and C131672, respectively. SR is defined as “Surgical removal of a part of a lesion; some portion of the lesion is detectable on post-operative evaluation.” Whereas GTR refers to “Surgical removal of an entire visible lesion, with no obvious lesion detected on post-operative evaluation; microscopic residual disease may be present.”. Follow-up records indicate that among the 53 GTR patients, fewer than 40% were later reported as tumor-free and did not experience cancer metastasis. Consequently, drug response assessment remains feasible according to RECIST criteria. The baseline patient clinical characteristic details can be found in Table S5.

### Only a few algorithm-omics pairs lead to predictive ML models

To harness the potential of machine learning algorithms and fairly evaluate the contribution of different omics data to predicting drug response, we assessed two distinct encoding strategies for drug response labels. The first approach involves binarizing the original four levels of drug response into two discrete classes as responder and non-responder for classification tasks. We defined a responder as a patient who had complete response (CR), partial response (PR) or stable disease (SD), and a non-responder as a patient who had progressive disease (PD). The second strategy entails encoding the four response levels as continuous numerical values, applicable to regression tasks. For the classification task, we evaluated models using both default hyperparameters and tuned hyperparameters. For regression, we evaluate with tuning hyperparameters only since hyperparameter-tuning shows a more stable performance evaluation in classification tasks. To be able to compare the regression performance with the performance of the classifier, we binarized the prediction of the regressor model with the threshold that maximises the MCC in the training set (Figure 1C). The performance of six candidate algorithms and their OMC variants was evaluated for each dataset. The effectiveness of each classifier exhibited variability contingent upon the data type, with miRNA, isomiR, and methylation consistently producing higher mMCC values compared to other data types, particularly when utilizing XGBoost with miRNA data. Conversely, classifiers such as CART and SVM demonstrated inferior performance across most data types. The analysis revealed that CNV data generally resulted in suboptimal classifier performance, with mMCC values ranging from -0.043 to 0.171. Notably, Methylation-RF demonstrated superior efficacy in both classification and regression tasks. Among the algorithms assessed, the XGB model based on miRNA data achieved the highest performance, with a mMCC of 0.447 (Figure 2A).

**Figure 2.**
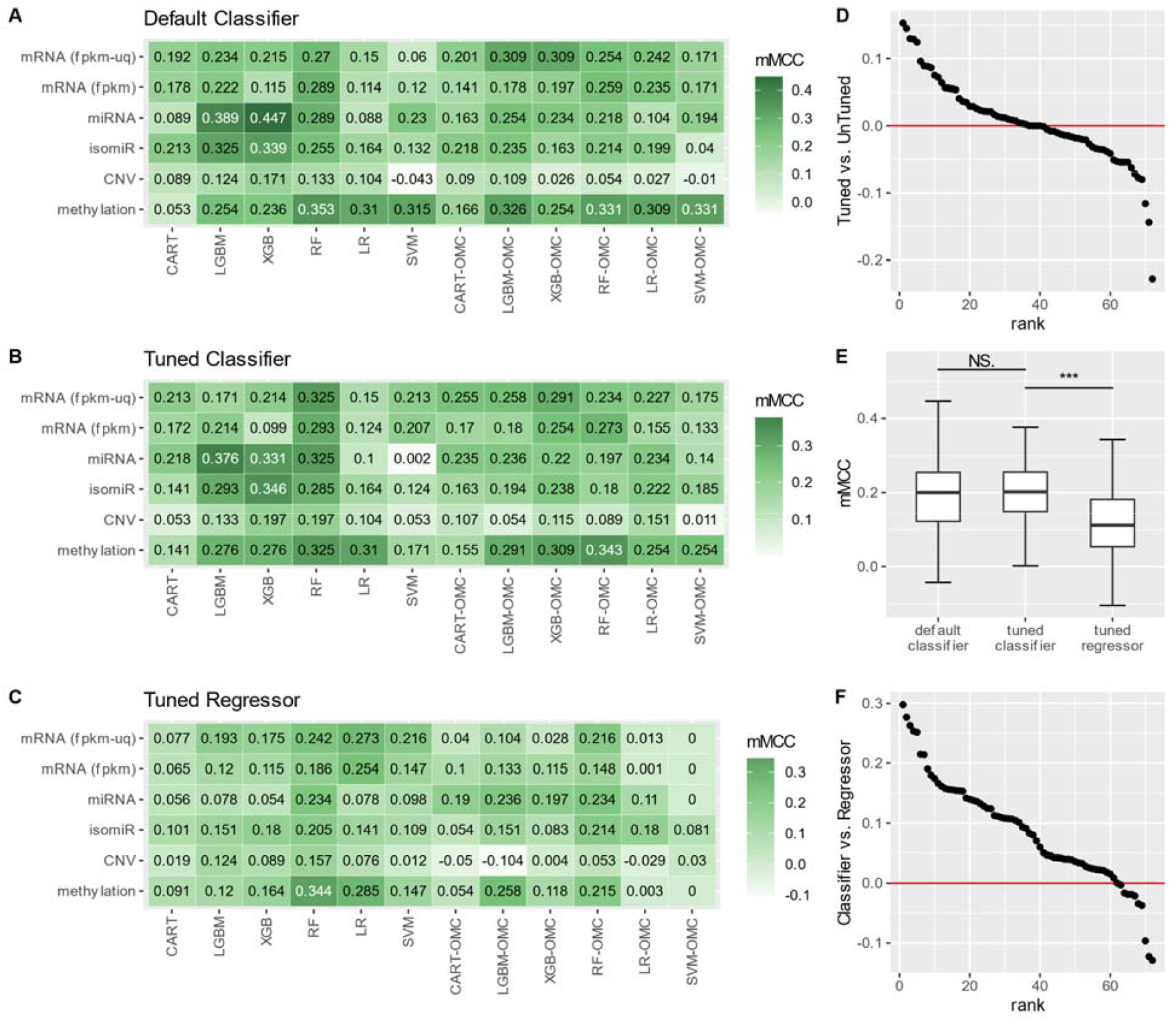
Out-of-sample performance comparison between regressors and classifiers with and without hyperparameter tuning. **A-C** 5-runs-median MCC of A classifiers with default hyperparameters, **B** classifiers with Bayesian hyperparameter tuning and **C** regressor with Bayesian hyperparameter tuning. Each block of the heatmap shows the median MCC of five evaluations of the predictive performance of each specific algorithm-omics pair. The y-axis shows the omics data used, and the x-axis shows the ML algorithm used. Each heatmap contains 12 ML algorithms including Classification and Regression Tree (CART), Light Gradient-Boosting Machine (LGBM), eXtreme Gradient Boosting (XGB), Random Forest (RF), Logistic Regression (LR), and Support Vector Machine with radial basis function kernel (SVM) and their OMC version. **D** Rank plot of the median MCC difference between between classifiers with and without hyperparameters tuning where each point represents one of the 72-dataset-algorithm pair. **E** Box plot of the median MCC distribution of classifiers with default hyperparameters, classifiers with hyperparameters tuning and regressors with hyperparameters tuning. **F** Rank plot of the median MCC difference between tuned classifiers and tuned regressors where each point represents one of the 72 dataset-algorithm pair.

We applied Bayesian optimizer for hyperparameter tuning^44^. The application of hyperparameter tuning through the Bayes optimizer enhances the median MCC of some models. However, it also results in a decrease in performance for certain models that initially exhibited favourable outcomes with default parameters (Figure 2B). Similarly, there is no significant difference between the overall median ROC-AUC and PR-AUC when hyperparameter tuning is performed (Figure S2). Thus, it is difficult to say whether the hyperparameter tuning procedure is good or bad for this task.

To compare the general performance of the classifier with and without hyperparameter tuning and the regressor with hyperparameter tuning, we compared the tuned and untuned model of each of the 72 data-algorithm pairs (6 data type × 12 algorithms). The two-sided paired Wilcoxon test is performed (n=72), revealing no statistically significant difference between the performance of classifiers using default hyperparameters and those with tuning (p-value = 0.45). However, the classifier outperforms the regressor significantly (p-value = 4.5×10^-10^). In contrast to classification tasks, regression tasks generally exhibit a decline in model efficacy, with the Random Forest (RF) model being a notable exception. This observation implies that RF may more effectively extract relevant information in regression contexts and handle non-linear feature relationships. Additionally, it can be recognised from Figure 2, that the OMC strategy tends to impair the performance of most models. This degradation could be attributable to a high false positive rate in feature selection or the ANOVA F-value’s limitations in capturing non-linear associations between features and outcomes.

### Comparing the performances of the systematic ML analysis and the auto-ML method JADBio

To expand our analysis, we use Just Add Data (JADBio), an auto-ML pipeline, to evaluate the same datasets each with 109 LGG patients with a history of temozolomide usage^45^. Instead of the NCV in the systematic ML analyses, JADBio introduced Bootstrap Bias Corrected (BBC)-CV to evaluate model performance. To permit a direct comparison, we partitioned the out-of-sample predictions of our model into five groups, each corresponding to a distinct random seed. BBC was applied to the models within each group, yielding the BBC-MCC. The median BBC-MCC across these five groups was then compared with the median value obtained from JADBio. The implementation of the BBC procedure is described in the Methods section.

The comparison shows that the performance of the systematic analysis is better than JADBio’s model in most omics data, but the performance is slightly lower than JADBio’s model when using the mRNA(fpkm-uq) profile or the isomiR profile.

**Table 1.**
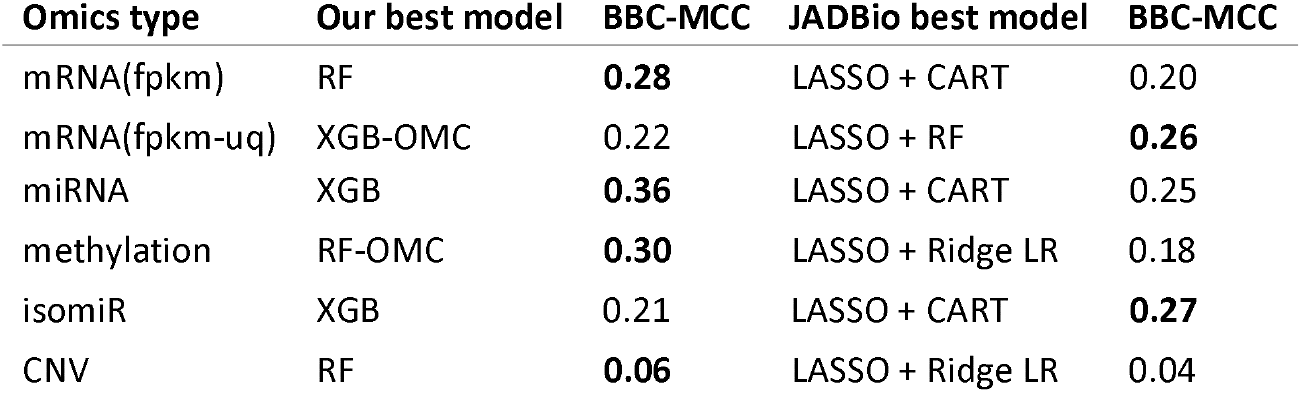
Compare the median MCC evaluated by NCV of the best classification model from our pipeline with the MCC evaluated by BBC-CV of the best model from JADBio. LASSO stands for Least Absolute Shrinkage and Selection Operator. For each data type, the best performance is highlighted in bold font.

| Omics type | Our best model | BBC-MCC | JADBio best model | BBC-MCC |
| --- | --- | --- | --- | --- |
| mRNA(fpkm) | RF | **0.28** | LASSO + CART | 0.20 |
| mRNA(fpkm-uq) | XGB-OMC | 0.22 | LASSO + RF | **0.26** |
| miRNA | XGB | **0.36** | LASSO + CART | 0.25 |
| methylation | RF-OMC | **0.30** | LASSO + Ridge LR | 0.18 |
| isomiR | XGB | 0.21 | LASSO + CART | **0.27** |
| CNV | RF | **0.06** | LASSO + Ridge LR | 0.04 |

### ML models based on miRNA profiles are more predictive than those based on MGMT profiles and especially than MGMT markers

MGMT methylation status is popular as a biomarker for predicting temozolomide response in gliomas^8^. The MGMT enzyme repairs temozolomide-induced DNA damage in tumor cells, thereby contributing to drug resistance. Hypermethylation of gene promoters is expected to lead to reduced gene expression by inhibiting access of transcription factors and the transcription machinery to DNA. Thus, the methylation status of the MGMT promoter serves as an indicator of favourable prognosis and response to temozolomide treatment in LGG. In clinical practice, various techniques can be employed to quantify the degree of methylation of MGMT, including Pyrosequencing (PSQ), quantitative methylation-specific PCR (qMSP), and immunohistochemistry (IHC). These techniques are used to measure only a small fraction of the methylation status in a 726bp-long CpG island (CGI) located on the promoter of the MGMT gene. The 70th-90th CpG among the 98 CpG sites on the CpG island is commonly used as the main indicator of MGMT promoter methylation status. These CpG sites are located on the island’s second differentially methylated region (DMR2), which plays a key role in controlling the MGMT promoter^46,47^.

In this study, we evaluated the predictive ability of MGMT methylation status for temozolomide response and compared it with the best miRNA-XGB model. In the clinical approach, the CpG sites and the cut-off for determining the methylation status (positive or negative) are highly depends on the technology. To reduce the impact of threshold selection, we directly use the real-valued MGMT DMR2 methylation beta value and the predicted probability to be responder of our model as the likelihood to be responder (Figure 3C).

**Figure 3.**
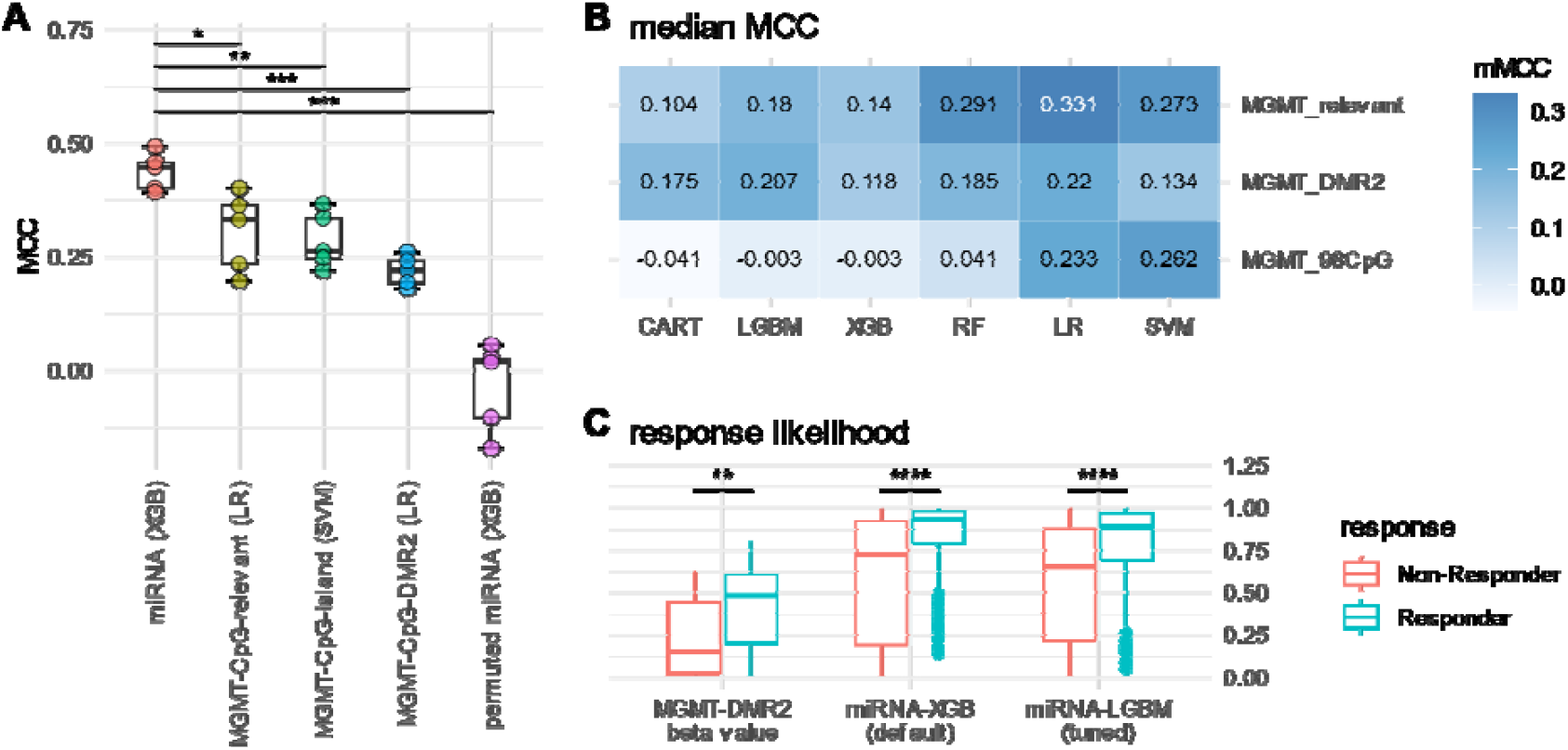
Comparison of the predictive performances of the best model to the baseline of MGMT and permuted model. **A** The significance level between the performance of our best model and baseline model. The labels on the x-axis indicate the dataset and ML algorithm. Each dot represents a run (of five random seeds). The significance between the best model and baseline is calculated by t-test. **B** The performance of models that predict drug response from MGMT relevant CpG beta value. **C** Comparison of the differences in likelihood between Responder and Non-Responder given by the biomarker MGMT-DMR2 and the ML model (Wilcoxon rank-sum test, two-sided. P-value = 1.03×10^-3^ for biomarker; p-value = 1.83×10^-13^ and p-value = 2.68×10^-12^ for ML with default and tuning hyperparameters).

To enhance the utility of biomarkers, we selected CpG sites within the MGMT methylation island, DMR2, and other MGMT-associated regions as features, resulting in three distinct datasets to assess the predictive efficacy of MGMT as a biomarker. The methodologies for probe localization on the MGMT methylation island and DMR2, as well as the extraction of MGMT-relevant CpG sites, are detailed in the methods section. Methylation beta values were obtained for 14 CpG sites within the methylation island and one site within DMR2. Subsequently, the predictive performance of these three MGMT-related datasets was evaluated using our analytical workflow. OMC feature selection strategy is not considered here due to the limited number of features.

The best ML model based on omics dataset has significantly better performance than the best MGMT-based ML model and especially than the MGMT markers (Figure 3). Comparison of the likelihood of the commonly used clinical biomarker MGMT-DMR2 with that of the ML model in the responder and non-responder showed that latter is significantly more predictive the former (Figure 3).

### Six predictive differential expressed miRNAs extracted from the best model

To identify predictive factors, we assessed the feature importance based on gain values of each variable within the optimal model (miRNA-XGB). We then retained the 30 features of the highest importance. Differential expression analysis of these 30 miRNAs was conducted using a two-sided Wilcoxon rank-sum test, followed by Bonferroni correction, resulting in the selection of 6 miRNAs with statistically significant differential expression (n=109).

As shown in Figure 4, the six differential expression miRNAs: hsa-miR-7-2, hsa-miR-6513, hsa-miR-376c, hsa-miR-335, hsa-miR-204, and hsa-let-7f-1, were found to be associated with resistance to temozolomide. hsa-miR-335 has been indicated by multiple studies that its high expression level contributes to shorter survival of glioma patients^48-50^. Our results also confirmed that a higher expression of miR-335 is associated with a less sensitive drug response. There is also evidence suggesting that hsa-miR-7 can inhibit the growth of gliomas through multiple pathways^51,52^. The precise role of hsa-miR-376c in gliomas cancers remains unclear. This microRNA exhibits tumor-suppressive effects in certain cancer types^53^, while showing a positive correlation with tumor progression in others^54-56^. The miR-204 is suggested to be a tumor suppressor factor that is able to enhance the sensitivity of temozolomide in glioblastoma (a type of HGG)^57-59^. However, in the TCGA-LGG dataset, miR-204 shows a negative correlation with the temozolomide sensitivity. This may suggest a different drug resistance mechanism between LGG and HGG. For let-7f and hsa-miR-6513, there is not sufficient evidence to determine their association with cancer progression or drug resistance.

**Figure 4.**
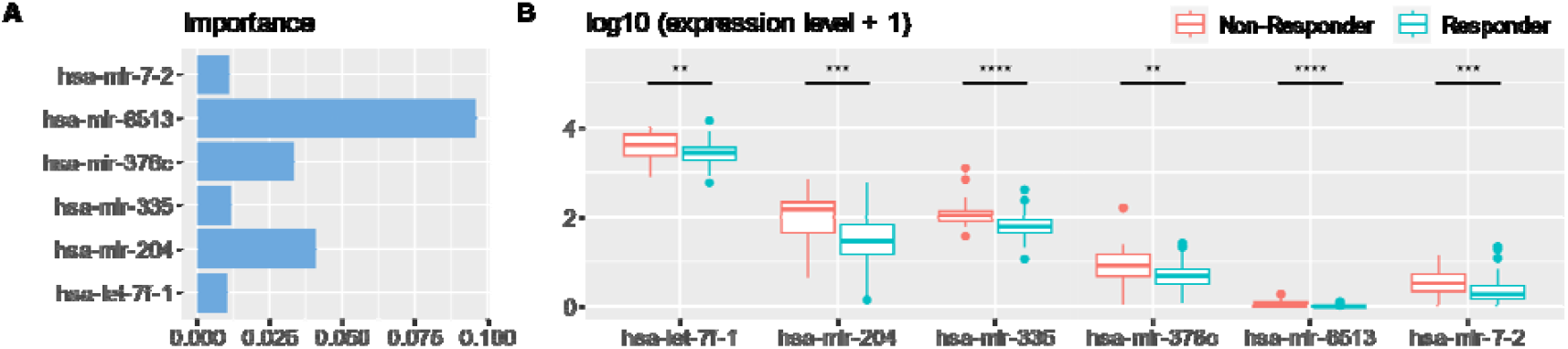
Feature importance analysis of the best model. The top 6 miRNA with highest feature importance in miRNA-XGB model. **B** The differentially expressed miRNAs among the top 6 miRNAs. The expression levels were added 1 and transformed into its logarithmic form for better visualization and comparison (log(x+1)). (Wilcoxon rank-sum test, two-sided. P-value = 3.39×10^-4^ for hsa-miR-7-2; P-value = 5.48×10^-5^ for hsa-miR-6513; P-value = 1.43×10^-3^ for hsa-miR-376c; P-value = 2.95×10^-5^ for hsa-miR-335; P-value = 4.88×10^-4^ for hsa-miR-204; P-value = 1.58×10^-3^ for hsa-let-7f-1)

### Ensemble models merging MGMT and miRNA data are only slightly more predictive

Considering that temozolomide resistance is influenced by various mechanisms beyond MGMT gene repair, we investigated whether integrating MGMT status with miRNA expression data enhances drug response prediction. Rather than directly combining miRNA and MGMT features into a single model, we employed an ensemble averaging approach with equal weight. This method facilitates a more equitable utilization of both datasets in predictive modelling. To be precise, we ensembled the best model trained on omics data (miRNA-XGB with default hyperparameter) with each MGMT model (methylation beta values of MGMT DMR2, MGMT promotor CpG island and MGMT relevant), respectively. As a result, the performance of the ensemble model has been slightly improved, where the MCC increases from 0.447 to 0.458 and the ROC-AUC increases from 0.670 to 0.687. However, the ensemble with MGMT did not significantly improve model performance compared to the best model using only miRNA (Figure 5). Considering that miRNA expression data are typically high-dimensional and already capture a wide array of regulatory information, the insignificant improvement implies that miRNA expression data may already encapsulate the information provided by MGMT status. This overlap means that adding MGMT features to the model doesn’t contribute new, independent information, leading to minimal improvement.

**Figure 5.**
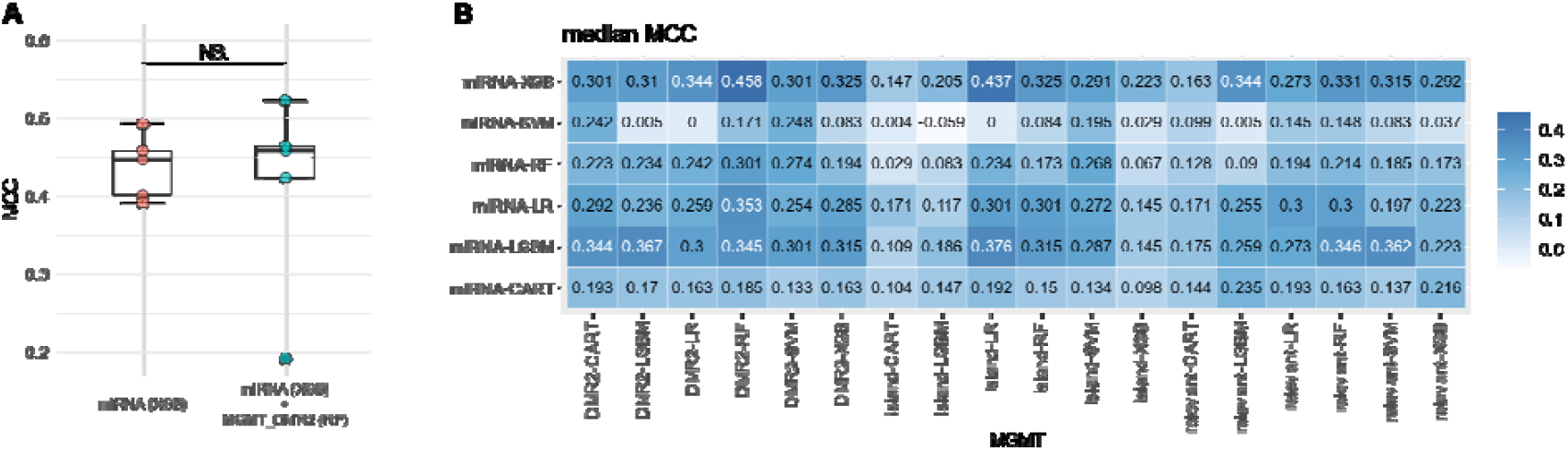
Performance of ensemble models. **A** Compare the best model performance between using omics data (miRNA-XGB) with the ensemble model of miRNA-XGB and the best model using MGMT (DMR2-RF). **B** The overall performance of all possible combinations of the miRNA models and three MGMT models (DMR2, CpG island, all relevant CpG sites).

### Adding clinical as features slightly improve model performance

Integrating clinical variables may bring non-redundant information to the model and thus improve the performance. Therefore, we evaluated the predictive power of 5 clinical variables: patient age, days between the biospecimen procurement and therapy start, cancer grade, procurement type and Karnofsky score. Patient age provide information about the overall physical health. The time interval between biospecimen collection and the initiation of therapy is critical for ensuring the integrity and viability of the samples used for diagnostic and predictive analyses. Additionally, cancer grade provides insight into the aggressiveness and differentiation of tumor cells. The Karnofsky Grade, which assesses the ability of the patient to perform daily activities, may reflect the overall quality of life and the impact of the disease.

To evaluate the linear association between these clinical variables and drug sensitivity, we utilized logistic regression models for each variable to predict drug response. The findings indicate that even when all five clinical variables are collectively employed as predictors, the Matthews correlation coefficient (MCC) remains marginal at 0.02, underscoring their limited predictive efficacy when considered independently (Table S4). Here, we are interested in quantifying the general improvement that adding clinical features could bring (to all models). As shown in Figure 2E, tuned and not-tuned models have the same median mMCC, and the tuned classifier has relatively less variation in performance (which means it is more stable). Such stability can better help us observe the improvement brought by clinical records to all models rather than just the best model. Therefore, the tuned model with a more stable performance is used for comparison instead of the default one.

We added all five clinical variables to the dataset as complementary features and evaluated them. Models that incorporate clinical data show higher mMCC compared to omics-only data, suggesting the added value of clinical information in enhancing predictive performance (Figure **6**). After adding clinical data, the MCC of the best model increased from 0.376 to 0.482, with methylation + clinical data and the LR-OMC model achieving the highest mMCC (0.482).

**Figure 6.**
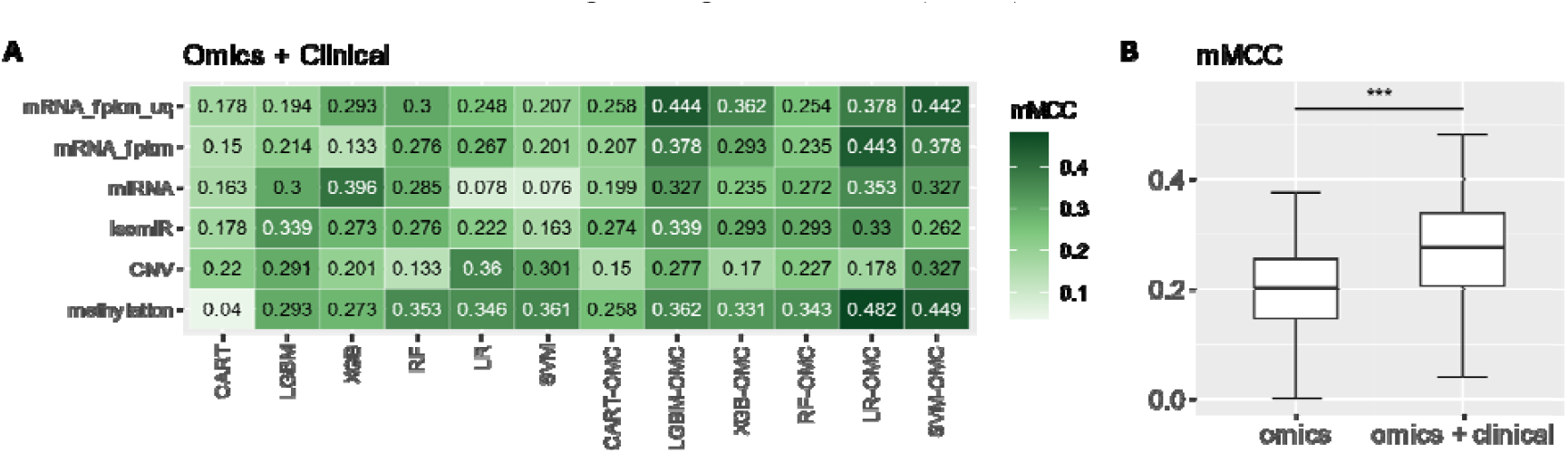
Model performance of classifier that train with additional clinical features. **A** Median MCC of each dataset-algorithm pair; **B** Comparison between models based on omics data only and models based on omics data and clinical variables as additional features (paired Wilcoxon rank-sum test, n=72, two-sided. P-value = 1.50×10^-7^).

### Bootstrap bias correction exposes the overestimation that may lead to the wrong conclusion

Considering that we evaluated many model-dataset combinations and looked for the best-performed model, even with a repeating of 5 with different random seeds and selected median performance, there is a risk of overestimating the best model performance. Suggesting we are evaluating n models, with the MCC performance { }, respectively. The distribution of the median MCC of the best-performed model can be represented as:

Suggest k is the best model and its performance is the performance we would like to estimate. Note that, for any value of x, the probability should be equal or smaller than 1. By retaining only, the evaluated performance and the theoretical best model performance, we have:

By flipping the sign, it can be shown that the probability of the evaluated best performance () to have a value higher than is never lower than the probability of the actual best model to have an MCC () higher than, thus inducing the risk of overestimation:

As an alternative to an unbiased estimation of the performance, we perform Bootstrap Bias Correction on out-of-sample prediction of all models we have evaluated^60^.

For example, to evaluate the unbiased performance of the best model shown in Figure 2A, that is, the unbiased performance of miRNA-XGB model, we need to take the out-of-sample predictions from all the models we have evaluated in Figure 2A and carry out the biased corrected process. Since in each run of evaluation (one random seed) there are 6 datasets and 12 algorithms that have been evaluated, the total sets of out-of-sample predictions should be . Because we evaluated 5 times with different random seeds, we need to perform bias correction on the results of each run, to obtain a total of 5 BBC-MCCs. The detail of the implementation of Bootstrap Bias Correction is explained in methods.

As shown in Figure 7, all bias-corrected performance (boxplot) exhibits a significant decline compared to the median MCC (diamond shape). Even so, the best models using omics data are still better than those using a single biomarker. Some conclusions can be influenced when comparing the bias-corrected performance with the original performance evaluated as median MCC. First, although the estimated MCC of the best model miRNA-XGB using the default hyperparameter was much higher than that of the best model miRNA-LGBM with hyperparameter tuning (median NCV-MCC 0.447 vs. 0.376), its median BBC-MCC of the five runs is very similar (median BBC-MCC 0.280 vs. 0.283), considering that the performance of the tuned model is more stable. Furthermore, our previous evaluation suggests that ensemble omics and MGMT could not significantly improve the performance, while the bias-corrected results indicated that the ensemble model has higher performance compared to its sub-model miRNA-XGB (median BBC-MCC 0.306 vs. 0.280).

**Figure 7.**
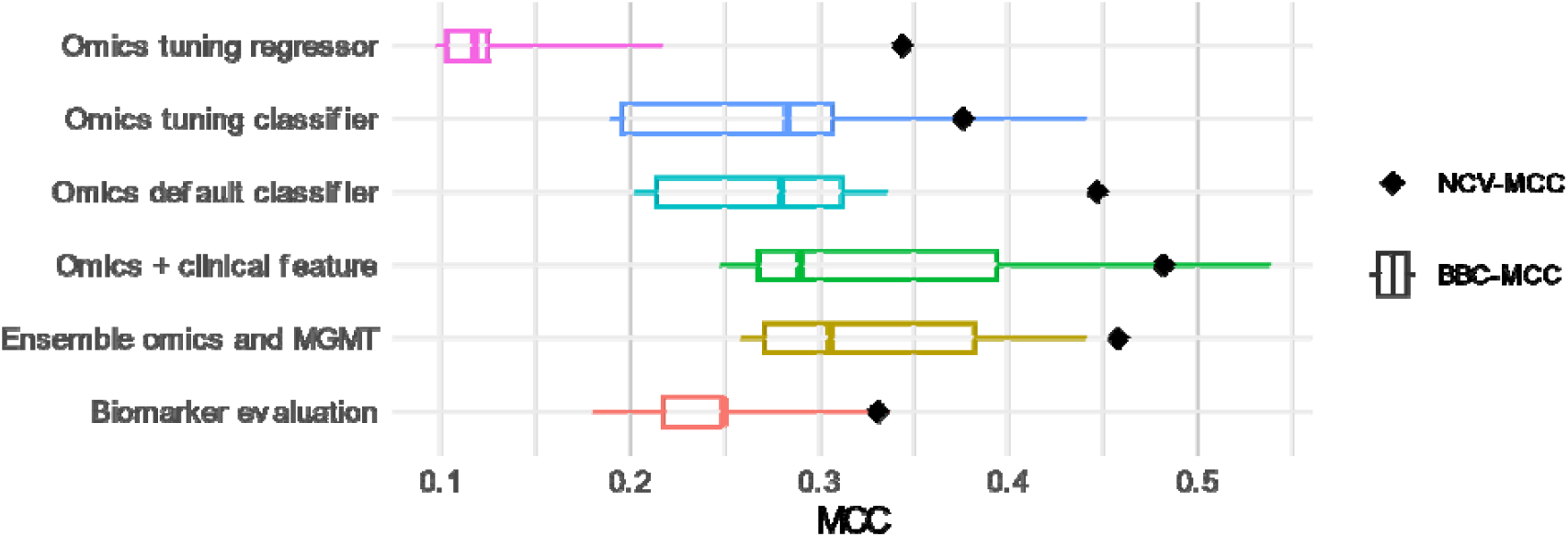
Comparison of unbiased performance of best model with different dataset. The boxplot shows the BBC-MCC of each run (with a specific random seed) and the diamond-shape point represent the median MCC evaluated by NCV.

## DISCUSSION

In this study, we aimed to evaluate the ability of supervised learning models to predict temozolomide drug response. For this, we utilized the high-quality dataset, which includes a broad type of omics data of 109 LGG patients with known temozolomide response. A wide range of machine learning algorithms is included in this study to identify the most predictive algorithm-omics combinations. Notably, the miRNA-XGB model exhibited superior predictive accuracy compared to models trained solely on the MGMT biomarker (MCC 0.447 vs. 0.331; ROC-AUC 0.710 vs. 0.693). Subsequent experiments revealed a notable enhancement in model performance upon incorporating specific clinical variables (from MCC 0.447 to 0.482), implying that clinical data may augment omics information to refine drug response predictions. These results indicate that the use of appropriate omics data and ML algorithms can reliably predict the drug response of LGG patients to temozolomide, thereby facilitating the selection of efficacious treatments and mitigating the risk of unnecessary chemotherapy-induced adverse effects.

Most predictive models for drug response are predominantly trained on preclinical datasets, notably cell-line data, rather than clinical tumor samples from patients^61^. This preference arises primarily because preclinical data, in contrast to clinical data, are more readily accessible, offer larger sample sizes, and provide more precise yet less clinically relevant labels, such as the Half-maximal inhibitory concentration (IC50). However, cell-line models are susceptible to various biases, including the absence of a tumor microenvironment, genetic drift, and selection biases inherent in the establishment of cell lines^62,63^. Consequently, cell-line-based models fail to capture the heterogeneous nature of clinical tumors, leading to suboptimal performance when applied to clinical data and thus are unsuitable for direct clinical deployment. This study exclusively utilized clinical data from patient samples, thereby enhancing clinical generalizability and demonstrating that, even with limited sample sizes, a significant level of predictive accuracy can be attained through the application of suitable machine-learning algorithms.

We identified several potential miRNA biomarkers for temozolomide drug sensitivity by applying feature importance analysis to the best-performed model. These include miR-335^48-50^ and miR-7^51,52^, which have been experimentally validated to be associated with glioma development, and miR-376c, which has been pointed out to be associated with the progression of other types of cancer. In addition, except for let-7f and miR-6513, whose relationship with cancer progression or drug resistance is still unclear, miR-204 has been suggested to have a tumor suppressor effect^64^. However, in GDC data, high expression of this miRNA is associated with temozolomide resistance. It is worth considering to conducting relevant experiments for these three miRNAs to explore the specific mechanisms.

It should be noted that, the estimation of MGMT performance in this study may be higher than the actual performance in the clinic. In the TCGA data employed, the methylation beta values were derived using the Infinium HumanMethylation450 array, which interrogates over 10 CpG sites dispersed across the MGMT CpG island. In contrast, clinical test kits typically assess fewer than 10 contiguous sites, resulting in reduced coverage. This broader coverage in our study may have conferred a more comprehensive representation of methylation status. Moreover, the threshold for determining methylation positivity varies among clinical test kits, which complicates direct comparisons. Given the lack of a standard reference from the HumanMethylation450 array, we selected three MGMT-related site sets—MGMT CpG island DMR2, MGMT CpG island, and MGMT-relevant sites for evaluation. The first two are commonly accessed in clinical practice, while the MGMT-relevant spot provided by the HumanMethylation450 manifest contains more than 100 sites, and its coverage far exceeds the scope of the CpG island of MGMT promoter. As an alternative to choosing an explicit cut-off, we used 12 ML algorithms to separately evaluate the ability of these three sets of MGMT datasets to predict drug response to maximize the potential of MGMT as a predictive factor. These factors may cause the estimated predictive power of MGMT to be higher than the actual clinical performance. Even so, our best model still showed significantly better performance than the baseline MGMT.

The difference between mMCC and BBC-MCC exposes the overestimation problem that can arise from evaluating multiple models on limited datasets. The overestimation can lead to high expectations for the performance of the best model and may cause unfair comparisons with other relevant studies. In our study, models utilizing default hyperparameters exhibit greater performance variance compared to classifiers with hyperparameter tuning, rendering them more susceptible to overestimation. This discrepancy is evident in the BBC-MCC metrics, where the top-performing default hyperparameter model, miRNA-XGB, displays a significantly lower BBC-MCC compared to its mMCC (median BBC-MCC = 0.280; median NCV-MCC = 0.447). Conversely, classifiers with tuned hyperparameters show a markedly smaller disparity between mMCC and BBC-MCC (median BBC-MCC = 0.283; median NCV-MCC = 0.376), indicating superior performance. However, reliance solely on the mMCC, which is prone to overestimation, obscures the enhancements provided by hyperparameter tuning and may mislead researchers into erroneously concluding that the default hyperparameter model performs better. Therefore, addressing overestimation is crucial in studies evaluating numerous models.

In this study, we analyzed data from 109 patients diagnosed with low-grade glioma (LGG), and our findings suggest that significant predictive accuracy can be achieved even within this limited sample size. However, to enhance the robustness and generalizability of our results, it would be advantageous to discuss the potential for including a larger dataset in future research. A more extensive dataset could provide greater statistical power and reduce the risk of overfitting, thereby allowing for more reliable predictions. Furthermore, considering that a large proportion of the 109 patients were from the United States, this may also impact our estimates of the model’s generalizability, as variations in demographic and clinical characteristics across different populations could influence the applicability of our findings. Incorporating an independent validation cohort would strengthen the validation of our predictive models, ensuring that they perform consistently across diverse populations. It is important to note that this analysis is conducted in silico, highlighting the necessity for experimental validation of our findings. Future studies should aim to corroborate our in-silico predictions through clinical trials or laboratory experiments, ultimately bridging the gap between computational analyses and practical applications in patient care.

## CONCLUSION

In conclusion, this study systematically evaluated the performance of multiple widely used models across diverse omics datasets, identifying models with outstanding predictive accuracy. Furthermore, the analysis provided valuable insights into the role of miRNA in temozolomide resistance. By integrating biomarkers and clinical data, we demonstrated a significant improvement in model performance, highlighting the value of incorporating multi-dimensional data. These findings contribute to a deeper understanding of the relationships between different datasets and algorithms, offering important direction for the future development of advanced artificial intelligence methods and their potential application in guiding treatment strategies for glioma patients.

## METHODS

### Molecular profiles acquisition from the TCGA dataset

The molecular profiles and clinical data of LGG patients were downloaded from the GDC portal [https://portal.gdc.cancer.gov]. The GDC provides the cancer research community with a unified data repository that enables data sharing across cancer genomic studies in support of precision medicine^38^. All omics and clinical data analysed in this study are publicly available. The filters for downloading the clinical data and manifest file for each molecular profile are listed in Table S1.

### Clinical data and drug response acquisition

To begin with, we merge the three clinical data: (1) A drug report containing the details of the drug name, drug type, the start and end day of treatment, and patient responses; (2) A specimen report containing sample procurement dates, method of sample procurement, and relevant information; (3) A clinical record containing age, tumor grade, and performance scale of the patient’s overall health status. These clinical data were integrated by matching records by patient ID (e.g. TCGA-3C-AALI). We then standardised the drug name. In the TCGA dataset, a given drug or treatment may have multiple names; therefore, standardising drug names is essential to maintain consistency in patient-drug pairs. The drug names were standardised using the drug correction table for the TCGA project provided by Spainhour et al.^65^.We filtered the clinical data to retain only patients with valid drug records, including drug name, drug response, therapy start date, and tumor procurement date. Patients who received drug treatments before tumor resections were excluded from the study. This is because the drugs might have affected and altered their tumor specimens, making them unsuitable to use as baseline samples. Lastly, some patients may have undergone the same treatment during multiple periods, leading to duplicate patient-drug records with different therapy start/end dates and multiple drug response records. In such cases, the consistency of the drug response record should be checked so that patient-drug pair with inconsistent drug response records can be removed.

### Data pre-processing for ML

To prepare the dataset for ML, we merged the encoded drug response from clinical data with each of the molecular data using the patient barcode as the key. For example, the molecular profile with barcode TCGA-DU-6397-02A-12R-A36C-13 corresponds to the clinical record of patient TCGA-DU-6397, as the first 10 characters represent the patient identifier. The input dataset is a two-dimensional matrix where each row corresponds to a patient sample and each column corresponds to a feature (e.g., gene expression levels, clinical variables). Then, it is necessary to ensure that there are no missing values in the matrix. We calculated the proportion of missing values for each feature. There are no missing values in both the mRNA and miRNA profiles. Since the missing value ratio of most features is very close to 0Thus, columns with any missing value are dropped. For patients with multiple samples, only the first sample (based on the alphabetical order of sample ID) is retained to avoid introducing bias into the data. For the mRNA profile, genes are filtered to retain only protein-coding genes. The number of samples and features after preprocessing is summarised in Table S3.

The drug response labels are encoded using two distinct strategies. For the classification task, complete response (CR), partial response (PR), and stable disease (SD) are encoded as 1, indicating a responder, while progressive disease (PD) is encoded as 0, indicating a non-responder. In the regression task, CR, PR, SD, and PD are assigned integer values from 0 to 3, respectively, representing varying degrees of drug resistance.

### Clinical feature encoding

The Karnofsky performance status scale, ranging from 0 to 100, was assessed both at the initial diagnosis and during the follow-up survey. Due to disease progression, in most cases, the two Karnofsky scores are inconsistent. Since our model aims to predict drug response for patients who have not yet taken medication, the follow-up data would not be available to us. Therefore, we only consider Karnofsky score from the primary diagnosis. 27.5% of the Karnofsky grades were missing and were imputed with the mean values of 87.2. Cancer grades II and III were encoded as 2 and 3, respectively. Patient age was recorded as at diagnosis. Procurement type was encoded as 0, 0.5, and 1, representing gross total procurement, subtotal procurement, and no procurement, respectively. Dates of procurement and therapy initiation were extracted from drug usage records and encoded in days. The baseline patient clinical characteristic details can be found in Table S5.

### Machine learning algorithm and OMC feature selection

All machine learning experiments were conducted in Python 3.11.0. Specifically, LR, SVM, CART, and RF algorithms were implemented using scikit-learn 1.2.0; LGBM was implemented using lightgbm 3.3.3; XGBoost was implemented using xgboost 1.7.1; The Bayes optimizer was implemented using scikit-optimize 0.9.0. All models were trained on high-performance computing clusters maintained by the Research Computing Service (RCS) team at Imperial College London.

The Optimal Model Complexity (OMC) search strategy was employed to reduce the feature and avoid overfitting. Specifically, we calculate the ANOVA F-value in the OMC search to rank all the features and then retain the top k features, where k ranges from 2 to n/2 (with n being the total number of features). The optimal value of k was evaluated in the inner loop of cross-validation. This method effectively reduces the number of features, the risk of overfitting, and the computational cost of model training.

To integrate the number of OMC features as a hyperparameter for tuning alongside other model parameters, we extended six algorithms (CART, LGBM, XGB, RF, LR, and SVM) by developing custom scikit-learn estimator APIs that accept the number of OMC features as an adjustable parameter in six derived child classes. Specifically, we overrode the parent classes, enabling the child classes to utilize only the top *k* features ranked by ANOVA F-value during both training and prediction, where *k* corresponds to the specified number of OMC features. Furthermore, the modified class accepts a feature list parameter, ensuring that any designated features are retained even if excluded from the OMC-selected subset. When additional clinical variables are included, all such variables are preserved in this manner.

### Bayesian optimization for hyperparameter tuning

Due to the significant computational demands associated with model evaluation, we employed the Bayes optimizer implemented in the Python package scikit-optimise (skopt) for hyperparameter tuning. The algorithm aims to fit a mapping function from hyperparameter to generalization error by the Gaussian process. Once the is obtained, the optimal hyperparameter can be obtained as the algorithm uses previous evaluation results to acquire more combinations of hyperparameters that are likely to be the optimal solution and evaluate these hyperparameter combinations in the next iteration. Therefore, it usually finds an acceptable combination of hyperparameters at a relatively lower computational cost. For classification and regression tasks, MCC and 1-R^2^ (where R^2^ is the coefficient of determination) are used as score functions to be maximised during hyperparameters tuning.

Hyperparameters were tuned in a 5-fold inner loop, and performance was evaluated in a 10-fold outer loop, forming a 5-fold-10-fold NCV framework. The hyperparameter combinations in each of the 10 outer loop iterations were usually different. According to the standard workflow of NCV, the performance of these 10 models is evaluated on the outer fold test set, which has not been used for training. This performance reflects how well the hyperparameter combinations perform, and the hyperparameter combination used by the best-performing model is treated as the optimal hyperparameter combination. However, due to the small size and imbalance of the data, the 10 sets of models are evaluated in one of the outer folds, which contain 11 samples at most. This results in colossal variance when evaluating the model directly on samples from a single fold. Moreover, considering that MMC is not a linear metric, its average may not have practical meaning. The model’s performance was not evaluated on the test set of each individual fold and subsequently averaged, as the size of these test sets would be too small, negatively impacting comparisons to cross-validation with other K values. To address this issue, a merged cross-validation approach was used. In this method, out-of-sample predictions from all folds were merged together, and the Matthews Correlation Coefficient (MCC) was calculated once using all 109 samples. Given that the dataset now includes four classes, with the smallest class (Partial Response) containing only five samples, we set the inner fold to 5 rather than 10 to improve the representation of all labels in each fold. The range of other hyperparameters are provided in Table S3.

### Extraction of MGMT methylation CpG sites

In the methylation profile collected from TCGA, the methylation beta values were obtained using the Illumina human methylation 450 BeadChip array. The probes corresponding to CpG sites relevant to MGMT were identified using the manifest provided by Illumina [available at support.illumina.com]. The position of each probe can be located by mapping the sites detected by each probe to the GRCh37 (human genome assemblies by the Genome Reference Consortium, build 37). By comparing them with the reference sequence of MGMT CpG island (CGI) located at position 131264948 - 131265710 on chromosome 10 (NCBI Gene ID: 4255), we identified probes within MGMT promoter CGI and the second differentially methylated regions (DMR2). As a result, we locate one probe on DMR2 (corresponding to the 84^th^ CpG site among sites 73-90^th^), 14 probes on the MGMT promoter CGI, and a total of 105 probes with no missing value relevant to MGMT (Figure 8).

**Figure 8.**
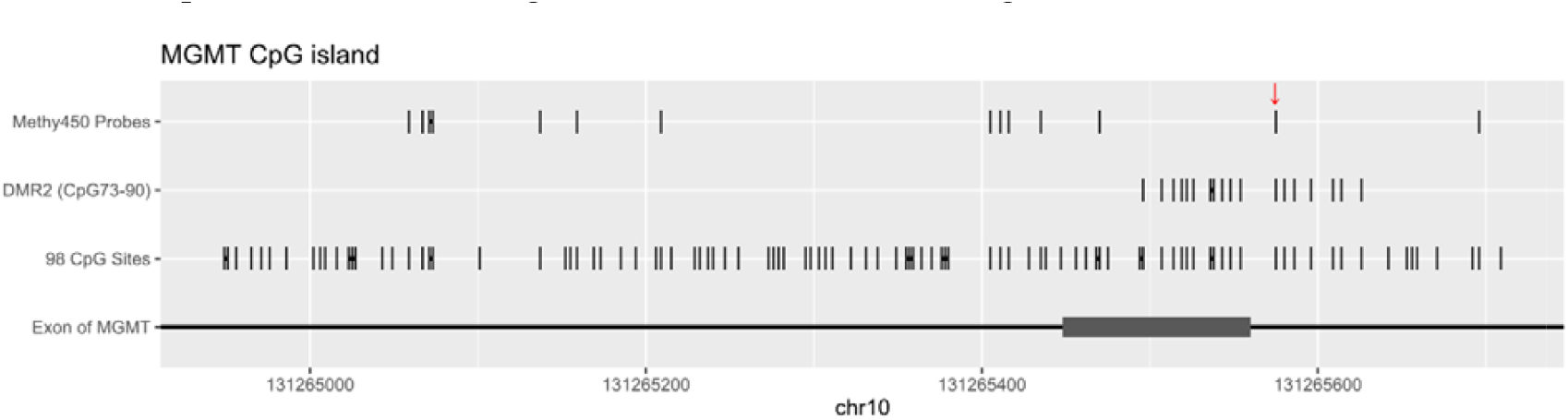
The CpG sites on the MGMT CGI. The figure shows 98 CpG sites located on the CGI, 14 sites on DMR2 (which are the 73-90^th^ CpG sites on CGI) and the CpG sites detected by Illumina human methylation 450. Adjacent CpG sites are connected by horizontal lines. Within the coverage region of the DMR2, only a solitary site was detected by the Illumina Human Methylation 450 assay, located at 131265575 with ID cg12981137, marked by a red arrow. In our study, the methylation beta value at this site is used to determine the methylation status.

### Model evaluation

We applied a 10-outer, 5-inner fold nested cross-validation (10-NCV) with multiple repetitions to mitigate the systematic overestimation of predictive performance that can result from selecting the best model among many candidate models. Using different random seeds to verify the same model repeatedly can significantly reduce the variance of performance estimation, thereby reducing such bias.

Given the imbalance in the dataset, MCC was chosen as the metrics for evaluating the model. MCC is a widely used as performance indicator in bioinformatics. It treats TP and TN equally, generating high scores only when both are high, making it reliable on imbalanced datasets. The predictions we e binarized using the threshold that maximised MCC on the training data. As a threshold-independent metric, the PR-AUC is considered to be better than ROC-AUC on imbalanced datasets. It has been noted that in the context of unbalanced data sets, ROC plots may seriously overestimate the reliability of the model, while PR-AUC is more robust. However, PR-AUC lacks many advantages of ROC-AUC, such as universal baseline, linear interpolation, optimality, and meaningful^66^. Therefore, in this study, we report both metrics.

### JADBio: autoML pipeline

We utilised the Auto-ML platform Just Add Data Bio (JADBio), accessible at www.JADBio.com^45^. JADBio aims to automate the sophisticated, advanced multivariate analysis of biological data and the discovery of biosignatures. JADBio handles the identification of ML problems, selection of candidate models, missing value imputation, feature selection, hyper-parameter tuning, performance evaluation and interpretation for the researchers. So far, JADBio offers a suite of ML algorithms for evaluation. For the classification task, the candidate models include Ridge Logistic Regression, Linear Support Vector Machine, Polynomial Support Vector Machine, Gaussian Support Vector Machine, Random Forest, and Decision Tree. For the regression task, the candidate models include Ridge Linear Regression, Random Forest, and Decision Tree.

JADBio employs 10-fold BBC-CV to estimate model performance. BBC-CV can reduce the systematic overestimation of its predictive performance due to selecting the best model from a considerable number of candidate models at a lower computational cost. Unlike the repeated NCV employed in our study, BBC-CV does not necessitate the retraining of models with identical configurations. Instead, it employs a bootstrap loop to ‘simulate’ the process of sample re-splitting thereby enhancing the accuracy of estimated performance at a low computational cost. Tsamardinos et al. quantitatively compared BBC-CV with general CV, suggesting that BBC-CV can reduce the estimation bias, but it has not been compared with repeated NCV.

### Bootstrap Bias Correction (BBC)

The MATLAB example code for BBC-CV is available on GitHub provided by Tsamardinos et al. at https://github.com/mensxmachina/BBC-CV^60^. Since this code was used to reproduce the simulation in their article and is not a standard BBC-CV implementation, we translated it into Python function. This function takes as input the out-of-sample prediction from all configurations (combinations of feature selection strategy, model training algorithm, and selected hyperparameter values), the true labels, and the number of BBC iterations, and outputs the bias-corrected performance (MCC) and the configuration of the best model. The detail steps and example of BBC-CV is provided in Figure S3.

## Availability of data and materials

The datasets supporting the results of this article are open-access data downloaded from the TCGA-LGG project within the GDC resource (https://portal.gdc.cancer.gov/projects/TCGA-LGG). The study reuses public data, and thus, no further ethical approval is required. The code is available in https://github.com/HanqinDu/TMZ-LGG

## Ethics approval and consent to participate

Not applicable as it is public data.

## Consent for publication

Not applicable as it is public data.

## Funding

This work was supported by the Commemoration of Her Royal Highness Princess Chulabhorn’s60th Birthday Anniversary scholarship from Chulabhorn Royal Academy (Thailand), the Marie-Curie actions IEF-Horizon (EU,LCII_PA4887), and the Wolfson Foundation and the Royal Society for a Royal Society Wolfson Fellowship (UK, RSWF\R1\221005).

## Authors’ contributions

P.J.B. and G.G. conceived the idea and designed the experiments. H.D. collected and analyzed the patients’ data, developed the ML process, and interpreted the results. H.D., C.P., G.G., and P.J.B. wrote the manuscript.

## Competing interests

The authors declare that they have no competing interests.

## Acknowledgements

We thank Yuanjie Zou for assistance in running the codes in the HPC facility and Zerui Li for assistance in generating plots, as well as their comments on early versions of the manuscript.

## Notes

### Competing Interest Statement

The authors have declared no competing interest.

## REFERENCES

1 Forst, D. A., Nahed, B. V., Loeffler, J. S. & Batchelor, T. T. Low-grade gliomas. The oncologist 19, 403–413 (2014).

2 Ostrom, Q. T. et al. The epidemiology of glioma in adults: a “state of the science” review. Neuro-Oncology 16, 896–913 (2014).

3 Ostrom, Q. T., Cioffi, G., Waite, K., Kruchko, C. & Barnholtz-Sloan, J. S. CBTRUS statistical report: primary brain and other central nervous system tumors diagnosed in the United States in 2014–2018. Neuro-Oncology 23, iii1–iii105 (2021).

4 Aldape, K. et al. Challenges to curing primary brain tumours. Nature reviews Clinical oncology 16, 509–520 (2019).

5 van Solinge, T. S., Nieland, L., Chiocca, E. A. & Broekman, M. L. D. Advances in local therapy for glioblastoma—taking the fight to the tumour. Nature Reviews Neurology 18, 221–236 (2022).

6 Denny, B. J., Wheelhouse, R. T., Stevens, M. F. G., Tsang, L. L. H. & Slack, J. A. NMR and molecular modeling investigation of the mechanism of activation of the antitumor drug temozolomide and its interaction with DNA. Biochemistry-Us 33, 9045–9051 (1994).

7 Rottenberg, D. A., Ginos, J. Z., Kearfott, K. J., Junck, L. & Bigner, D. D. In vivo measurement of regional brain tissue pH using positron emission tomography. Annals of Neurology: Official Journal of the American Neurological Association and the Child Neurology Society 15, 98–102 (1984).

8 Dunn, J. et al. Extent of MGMT promoter methylation correlates with outcome in glioblastomas given temozolomide and radiotherapy. British journal of cancer 101, 124–131 (2009).

9 Hegi, M. E. et al. MGMT gene silencing and benefit from temozolomide in glioblastoma. New Engl J Med 352, 997–1003 (2005).

10 Kontogeorgos, G. & Thodou, E. Is MGMT the best marker to predict response of temozolomide in aggressive pituitary tumors? Alternative markers and prospective treatment modalities. Hormones 18, 333–337 (2019).

11 Brandes, A. A. et al. O 6-methylguanine DNA-methyltransferase methylation status can change between first surgery for newly diagnosed glioblastoma and second surgery for recurrence: clinical implications. Neuro-Oncology 12, 283–288 (2010).

12 Amatu, A. et al. Tumor MGMT promoter hypermethylation changes over time limit temozolomide efficacy in a phase II trial for metastatic colorectal cancer. Ann Oncol 27, 1062–1067 (2016).

13 Liu, L., Markowitz, S. & Gerson, S. L. Mismatch repair mutations override alkyltransferase in conferring resistance to temozolomide but not to 1, 3-bis (2-chloroethyl) nitrosourea. Cancer Res 56, 5375–5379 (1996).

14 Karayan-Tapon, L. et al. Prognostic value of O 6-methylguanine-DNA methyltransferase status in glioblastoma patients, assessed by five different methods. Journal of neuro-oncology 97, 311–322 (2010).

15 Wood, R. D., Mitchell, M., Sgouros, J. & Lindahl, T. Human DNA repair genes. Science 291, 1284–1289 (2001).

16 Hanawalt, P. C. Subpathways of nucleotide excision repair and their regulation. Oncogene 21, 8949–8956 (2002).

17 Fu, D., Calvo, J. A. & Samson, L. D. Balancing repair and tolerance of DNA damage caused by alkylating agents. Nat Rev Cancer 12, 104–120 (2012).

18 Liu, B. et al. LncRNA SOX2OT promotes temozolomide resistance by elevating SOX2 expression via ALKBH5-mediated epigenetic regulation in glioblastoma. Cell Death Dis 11, 384 (2020).

19 Messaoudi, K., Clavreul, A. & Lagarce, F. Toward an effective strategy in glioblastoma treatment. Part I: resistance mechanisms and strategies to overcome resistance of glioblastoma to temozolomide. Drug Discov Today 20, 899–905 (2015).

20 Azuaje, F. Computational models for predicting drug responses in cancer research. Brief Bioinform 18, 820–829 (2017).

21 Costello, J. C. et al. A community effort to assess and improve drug sensitivity prediction algorithms. Nat Biotechnol 32, 1202–1212 (2014).

22 An, X., Chen, X., Yi, D., Li, H. & Guan, Y. Representation of molecules for drug response prediction. Brief Bioinform 23, bbab393 (2022).

23 Menden, M. P. et al. Machine learning prediction of cancer cell sensitivity to drugs based on genomic and chemical properties. Plos One 8, e61318 (2013).

24 Naulaerts, S., Menden, M. P. & Ballester, P. J. Concise polygenic models for cancer-specific identification of drug-sensitive tumors from their multi-omics profiles. Biomolecules 10, 963 (2020).

25 Li, H., Li, T., Quang, D. & Guan, Y. Network propagation predicts drug synergy in cancers. Cancer Res 78, 5446–5457 (2018).

26 Park, A., Lee, Y. & Nam, S. A performance evaluation of drug response prediction models for individual drugs. Sci Rep-Uk 13, 11911 (2023).

27 Baptista, D., Ferreira, P. G. & Rocha, M. Deep learning for drug response prediction in cancer. Brief Bioinform 22, 360–379 (2021).

28 Wilding, J. L. & Bodmer, W. F. Cancer cell lines for drug discovery and development. Cancer Res 74, 2377–2384 (2014).

29 Park, S., Soh, J. & Lee, H. Super. FELT: supervised feature extraction learning using triplet loss for drug response prediction with multi-omics data. BMC bioinformatics 22, 269 (2021).

30 Ertel, A., Verghese, A., Byers, S. W., Ochs, M. & Tozeren, A. Pathway-specific differences between tumor cell lines and normal and tumor tissue cells. Mol Cancer 5, 1–13 (2006).

31 Huang, E. W., Bhope, A., Lim, J., Sinha, S. & Emad, A. Tissue-guided LASSO for prediction of clinical drug response using preclinical samples. Plos Comput Biol 16, e1007607 (2020).

32 Jiang, G. et al. Comprehensive comparison of molecular portraits between cell lines and tumors in breast cancer. Bmc Genomics 17, 281–301 (2016).

33 Domcke, S., Sinha, R., Levine, D. A., Sander, C. & Schultz, N. Evaluating cell lines as tumour models by comparison of genomic profiles. Nat Commun 4, 2126 (2013).

34 Gillet, J.-P. et al. Redefining the relevance of established cancer cell lines to the study of mechanisms of clinical anti-cancer drug resistance. Proceedings of the National Academy of Sciences 108, 18708–18713 (2011).

35 Piyawajanusorn, C., Nguyen, L. C., Ghislat, G. & Ballester, P. J. A gentle introduction to understanding preclinical data for cancer pharmaco-omic modeling. Brief Bioinform 22, bbab312 (2021).

36 Jensen, M. A., Ferretti, V., Grossman, R. L. & Staudt, L. M. The NCI Genomic Data Commons as an engine for precision medicine. Blood, The Journal of the American Society of Hematology 130, 453–459 (2017).

37 Grossman, R. L. et al. Toward a shared vision for cancer genomic data. New Engl J Med 375, 1109–1112 (2016).

38 Weinstein, J. N. et al. The cancer genome atlas pan-cancer analysis project. Nat Genet 45, 1113–1120 (2013).

39 Ballester, P. J. & Carmona, J. Artificial intelligence for the next generation of precision oncology. Npj Precis Oncol 5, 79 (2021).

40 Bomane, A., Gonçalves, A. & Ballester, P. J. Paclitaxel response can be predicted with interpretable multi-variate classifiers exploiting DNA-methylation and miRNA data. Front Genet 10, 1041 (2019).

41 Ogunleye, A. Z., Piyawajanusorn, C., Gonçalves, A., Ghislat, G. & Ballester, P. J. Interpretable machine learning models to predict the resistance of breast cancer patients to doxorubicin from their microRNA profiles. Adv Sci 9, 2201501 (2022).

42 Dang, C. C., Peón, A. & Ballester, P. J. Unearthing new genomic markers of drug response by improved measurement of discriminative power. BMC Medical Genomics 11, 1–14 (2018).

43 Ogunleye, A., Piyawajanusorn, C., Ghislat, G. & Ballester, P. J. Large-Scale Machine Learning Analysis Reveals DNA Methylation and Gene Expression Response Signatures for Gemcitabine-Treated Pancreatic Cancer. Health Data Science 4, 0108 (2024).

44 Wu, J. et al. Hyperparameter optimization for machine learning models based on Bayesian optimization. Journal of Electronic Science and Technology 17, 26–40 (2019).

45 Tsamardinos, I. et al. Just Add Data: automated predictive modeling for knowledge discovery and feature selection. Npj Precis Oncol 6, 38 (2022).

46 Malley, D. S. et al. A distinct region of the MGMT CpG island critical for transcriptional regulation is preferentially methylated in glioblastoma cells and xenografts. Acta neuropathologica 121, 651–661 (2011).

47 Mansouri, A. et al. MGMT promoter methylation status testing to guide therapy for glioblastoma: refining the approach based on emerging evidence and current challenges. Neuro-Oncology 21, 167–178 (2019).

48 Teng, J.-W., Bian, S.-S., Kong, P. & Chen, Y.-G. Icariin triggers osteogenic differentiation of bone marrow stem cells by up-regulating miR-335–5p. Experimental Cell Research 414, 113085 (2022).

49 Wang, W. J. et al. LncRNA HANR aggravates the malignant progression of glioma via targeting miRNA-335. European Review for Medical & Pharmacological Sciences 24 (2020).

50 Luo, K. et al. CircKIF4A promotes glioma growth and temozolomide resistance by accelerating glycolysis. Cell Death Dis 13, 740 (2022).

51 Bao, X. a., Peng, Y., Shen, J. & Yang, L. Sevoflurane inhibits progression of glioma via regulating the HMMR antisense RNA 1/microRNA-7/cyclin dependent kinase 4 axis. Bioengineered 12, 7893–7906 (2021).

52 Pei, Y. et al. Circular RNA circRNA_0067934 promotes glioma development by modulating the microRNA miR-7/Wnt/β-catenin axis. Bioengineered 13, 5792–5802 (2022).

53 Iwaki, J. et al. MiR-376c down-regulation accelerates EGF-dependent migration by targeting GRB2 in the HuCCT1 human intrahepatic cholangiocarcinoma cell line. Plos One 8, e69496 (2013).

54 Cao, X., Zhang, J., Apaer, S., Yao, G. & Li, T. microRNA-19a-3p and microRNA-376c-3p promote hepatocellular carcinoma progression through SOX6-mediated Wnt/β-catenin signaling pathway. International Journal of General Medicine, 89–102 (2021).

55 McDonald, A. C. et al. Circulating microRNAs in plasma among men with low[grade and high[grade prostate cancer at prostate biopsy. The Prostate 79, 961–968 (2019).

56 Zheng, Y., Li, Z., Yang, S., Wang, Y. & Luan, Z. CircEXOC6B suppresses the proliferation and motility and sensitizes ovarian cancer cells to paclitaxel through miR-376c-3p/FOXO3 axis. Cancer Biotherapy & Radiopharmaceuticals 37, 802–814 (2022).

57 Li, T., Pan, H. & Li, R. The dual regulatory role of miR-204 in cancer. Tumor Biology 37, 11667–11677 (2016).

58 Yang, Y. N., Zhang, X. H., Wang, Y. M., Zhang, X. & Gu, Z. miR-204 reverses temozolomide resistance and inhibits cancer initiating cells phenotypes by degrading FAP-α in glioblastoma. Oncology Letters 15, 7563–7570 (2018).

59 Xia, Z., Liu, F., Zhang, J. & Liu, L. Decreased expression of MiRNA-204-5p contributes to glioma progression and promotes glioma cell growth, migration and invasion. Plos One 10, e0132399 (2015).

60 Tsamardinos, I., Greasidou, E. & Borboudakis, G. Bootstrapping the out-of-sample predictions for efficient and accurate cross-validation. Machine learning 107, 1895–1922 (2018).

61 Partin, A. et al. Deep learning methods for drug response prediction in cancer: predominant and emerging trends. Frontiers in medicine 10, 1086097 (2023).

62 Tseng, Y.-Y. & Boehm, J. S. From cell lines to living biosensors: new opportunities to prioritize cancer dependencies using ex vivo tumor cultures. Current Opinion in Genetics & Development 54, 33–40 (2019).

63 Hynds, R. E., Vladimirou, E. & Janes, S. M. Vol. 11 dmm037366 (The Company of Biologists Ltd, 2018).

64 Zhou, H., Ma, Y., Zhong, D. & Yang, L. Knockdown of lncRNA HOXD-AS1 suppresses proliferation, migration and invasion and enhances cisplatin sensitivity of glioma cells by sponging miR-204. Biomedicine & Pharmacotherapy 112, 108633 (2019).

65 Spainhour, J. C. G. & Qiu, P. Identification of gene-drug interactions that impact patient survival in TCGA. BMC bioinformatics 17, 1–8 (2016).

66 Flach, P. & Kull, M. Precision-recall-gain curves: PR analysis done right. Advances in neural information processing systems 28 (2015).

