## Supplementary file for "Predicting temozolomide response in low-grade glioma patients with large-scale machine learning"

Supplementary information

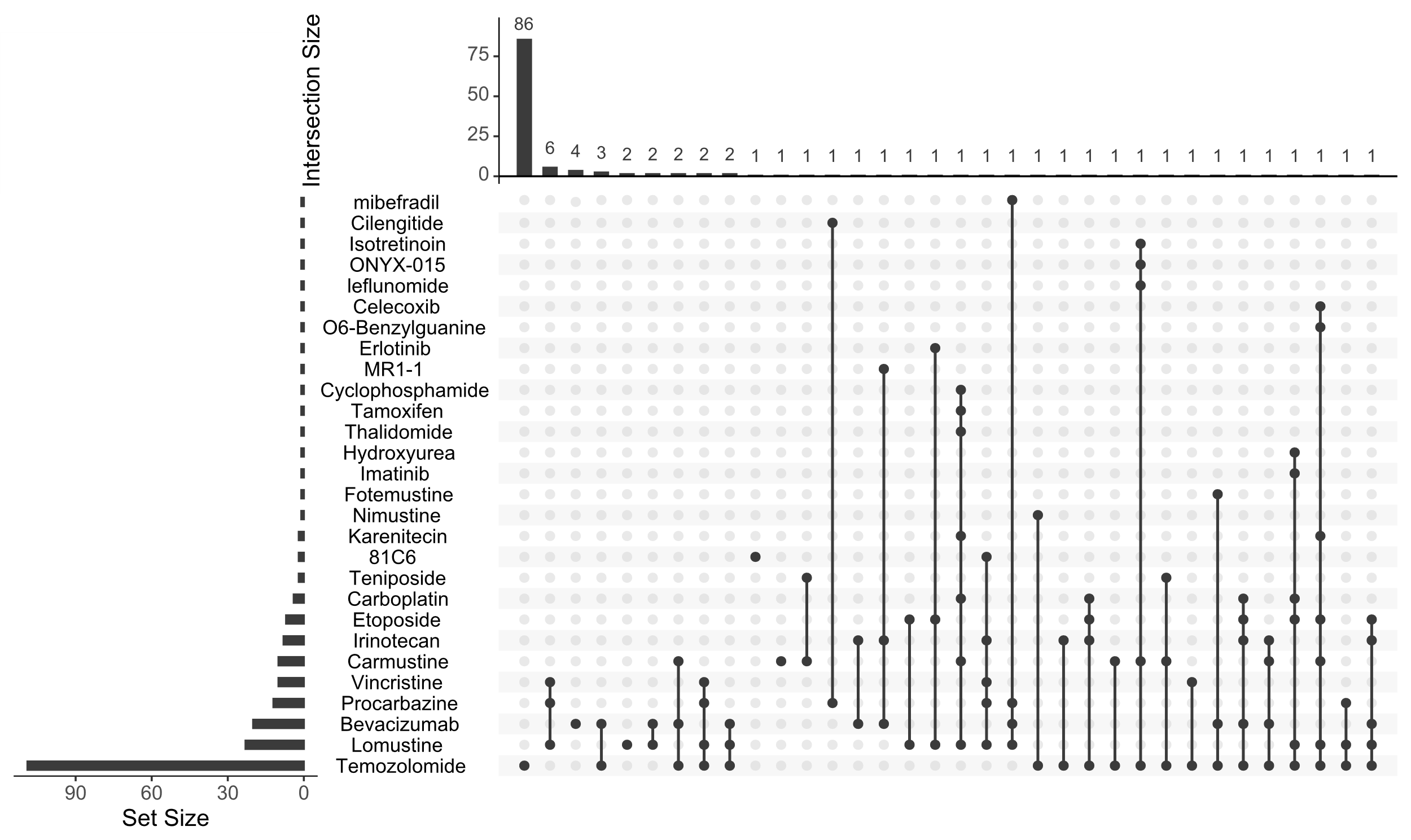

Figure S1. Of the 134 patients, nearly half of them had used more than one type of drug during the treatment. 86 of the patients have taken only temozolomide throughout the whole treatment. In most cases, patients take only one drug alone at a period. Of the 109 patients who had used temozolomide, only five patients had taken other medicines while taking temozolomide. Specifically, four patients took Bevacizumab (An antibody that blocks angiogenesis), and one of these patients also took Celecoxib (an anti-inflammatory drug). The last patient took Celecoxib (an anti-inflammatory drug). Since 86 of the 109 patients subject to temozolomide take only one drug alone at a period, the potential combined effect of drugs could be ignored.

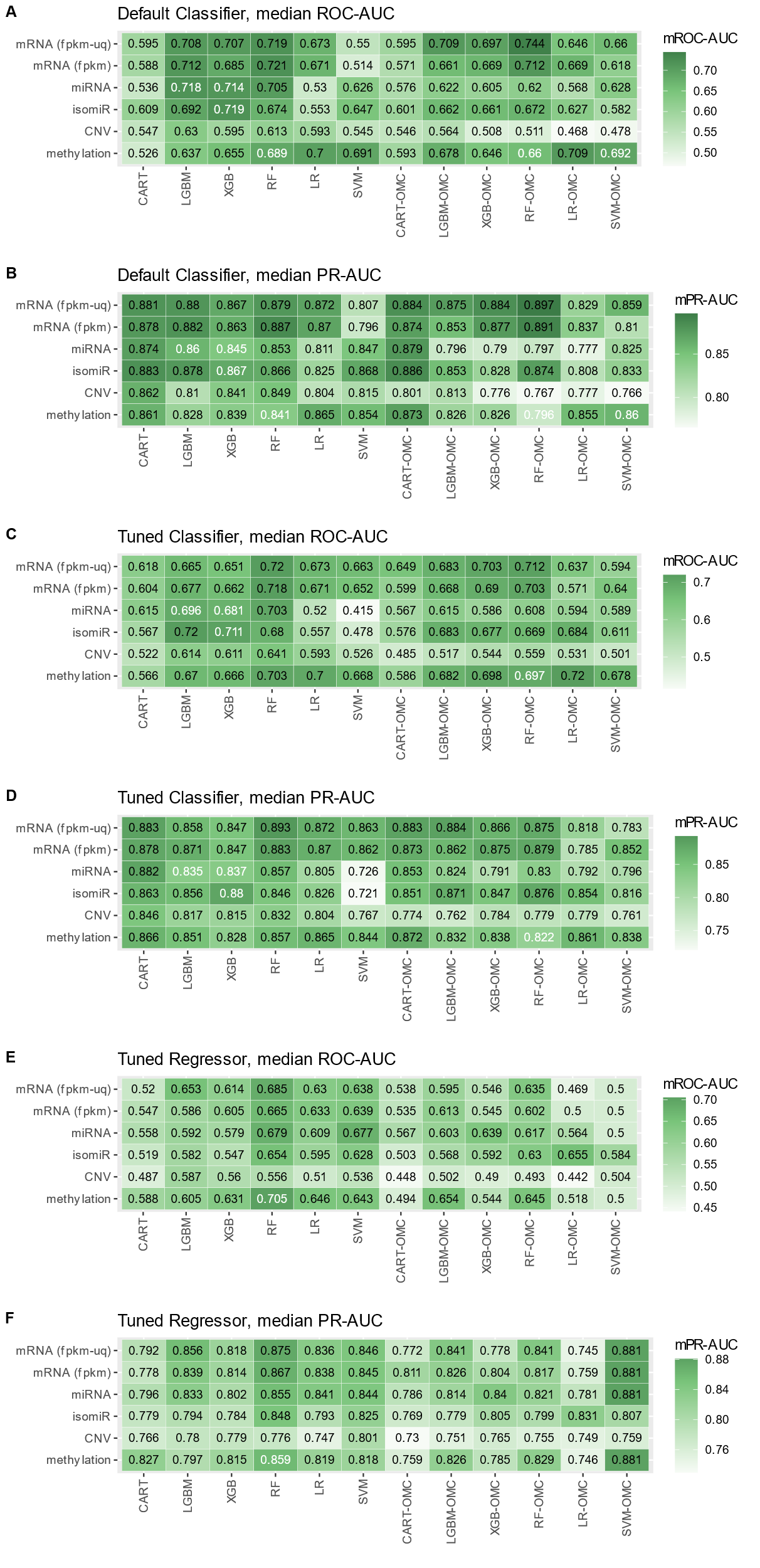

Figure S2. Comparison of the predictive ability of regressor and classifier with and without hyperparameter tuning. A-C Compares the median ROC-AUC and the D-F compares the median PR-AUC.

| Dataset | Filter |
| --- | --- |
| clinical record | General → Project = “TCGA-LGG”; Available Data → Data format = “bcr biotab” |
| mRNA | General → Project = “TCGA-LGG”; Available Data → Data Type = “Gene Expression Quantification”; Available Data →Workflow Type = “STAR – Counts”; |
| miRNA | General → Project = “TCGA-LGG”; Available Data → Data Type = “miRNA Expression Quantification”; |
| isomiR | General → Project = “TCGA-LGG”; Available Data → Data Type = “Isoform Expression Quantification”; |
| methylation β value | General → Project = “TCGA-LGG”; Available Data → Data Type = “Methylation Beta Value”; |
| CNV | General → Project = “TCGA-LGG”; Available Data → Data Type = “Gene Level Copy Number”; Available Data → Workflow Type = “ASCAT2”; |

Table S1. The filters to be applied for getting each dataset from GDC. The term may change with new version of GDC releases.

| Molecular profiles | Workflow Type | Patients treated by temozolomide (CR/PR/SD/PD) | Number of Features |
| --- | --- | --- | --- |
| mRNA (FPKM and FPKM-uq) | STAR - Counts | 109 (13/5/65/26) | 19962 |
| miRNA (RPM) | BCGSC miRNA Profiling | 109 (13/5/65/26) | 1881 |
| isomiR (RPM) | BCGSC miRNA Profiling | 109 (13/5/65/26) | 1149 |
| CNV | ASCAT2(Affymetrix SNP 6.0) | 109 (13/5/65/26) | 59881 |
| methylation beta value | SeSAMe Methylation Beta Estimation (Illumina Human Methylation 450) | 109 (13/5/65/26) | 371142 |

Table S2. Overview of the ML-ready dataset. CR, PR, SD and PD represent the number of "Complete Response", "Partial Response", "Stable Disease", and "Clinical Progressive Disease", respectively, which are judged by RECIST criteria. The mRNA expression level is quantified by mean and upper quartile of Reads per kilobase per million mapped reads (FPKM and FPKM_uq); the miRNA and isomiR are quantified by Reads per million mapped (RPM).

| Algorithm | Hyperprameters | Range |
| --- | --- | --- |
| CART | criterion | {'squared_error', 'absolute_error', 'poisson'} |
|  | max_depth | ℤ ∈ [2, p^0.25] |
|  | min_samples_split | ℤ ∈ [2, 10] |
|  | k_omc | ℤ ∈ [1, n/2] |
| RF | criterion | {'squared_error', 'absolute_error', 'poisson'} |
|  | max_features | ℝ ∈ [0.01, 1.0] |
|  | max_depth | ℤ ∈ [2, log(p)] |
|  | min_samples_leaf | ℤ ∈ [1, 5] |
|  | k_omc | ℤ ∈ [1, n/2] |
| XGB | grow_policy | {'depthwise', 'lossguide'} |
|  | base_score | ℝ ∈ [0.1, 0.9] |
|  | reg_alpha | ℝ ∈ [0.0, 3.0] |
|  | reg_lambda | ℝ ∈ [0.0, 3.0] |
|  | max_depth | ℤ ∈ [2, p^0.25] |
|  | max_leaves | ℤ ∈ [3, 2*p^0.25] |
|  | k_omc | ℤ ∈ [1, n/2] |
| LGBM | boosting_type | {'gbdt', 'dart', 'goss'} |
|  | min_child_weight | ℝ ∈ [0.0001, 0.01] |
|  | reg_lambda | ℝ ∈ [0.0, 3.0] |
|  | reg_lambda | ℝ ∈ [0.0, 3.0] |
|  | max_depth | ℤ ∈ [2, p^0.25] |
|  | num_leaves | ℤ ∈ [3, 2*p^0.25] |
|  | min_child_samples | ℤ ∈ [5, 40] |
|  | k_omc | ℤ ∈ [1, n/2] |
| LR | C | ℝ ∈ [0.1, 10.0] |
|  | k_omc | ℤ ∈ [1, n/2] |
| SVM | kernel | {'linear', 'poly', 'rbf', 'sigmoid'} |
|  | tol | ℝ ∈ [0.0001, 0.1] |
|  | epsilon | ℝ ∈ [0.5, 3.0] |
|  | k_omc | ℤ ∈ [1, n/2] |

Table S3. The range of hyperparameters for tuning. The n and p represent the number of samples and number of features of the dataset respectively.

| **Clinical record** | **ROC-AUC** | **MCC** | **MCC (threshold tuning)** | **PCC** |
| --- | --- | --- | --- | --- |
| Age | 0.5 | 0 | 0 | -0.05 |
| Cancer grade | 0.5 | 0 | 0 | 0.04 |
| Karnofsky grade  Procurement type  Treatment start date  All above clinical records | 0.5  0.5  0.5  0.5 | -0.05  0  0  0 | 0  0  0.04  0.02 | 0.15  0.07  -0.25  / |

Table S4. Performance of logistic regression model predicting drug response with only clinical record. PCC stand for Pearson correlation coefficient between the clinical feature and drug response. The last row aim to evaluate the predictive ability of all extracted clinical features. Since we only calculate PCC between feature and label, PCC of last row (which include multiple features) is not available.

| Variable | Responder (N=83) | Non-Responder (N=26) |
| --- | --- | --- |
| Age, years, mean (SD) | 43.2 (13.1) | 45.0 (15.8) |
| Days to therapy start, mean (SD) | 190 (561) | 409 (451) |
| Karnofsky score, mean (SD) | 87.3 (9.22) | 83.6 (13.1) |
| Procurement type |  |  |
| Gross Total Resection | 39 | 14 |
| Subtotal Resection | 43 | 12 |
| Biopsy | 1 | 0 |
| Tumor grade |  |  |
| Grade 2 | 22 | 8 |
| Grade 3 | 61 | 18 |

Table S5. baseline patient clinical characteristic table.

**The BBC can be described as following steps:**

1. Given a set of configurations (combination of feature selection strategy, model training algorithm and selected hyperparameter values), calculate the out-of-sample prediction $\Pi$ of each data instance by 10-fold CV for each configuration. Where each row of $\Pi$ represents a data instance and each column of $\Pi$ represents a specific configuration. For example, if we have 50 data instances and 10 candidate configurations, the shape of $\Pi$ would be $50\times10$ where $\pi_{ij}$ is the prediction on sample i from the model with configuration j.
2. Each BBC iteration consists of randomly picking (with replacement) N rows from $\Pi$, that is, the out-of-sample of N of the samples from all configurations. This step aims to bootstrap a new sample set that belongs to the same distribution from valid samples by bootstrapping. With the resampled data of the predicted value, calculate the metrics from the out-of-sample prediction $\pi$ and the truth value $y$ to select the best configuration and evaluate its performance on the non-picked data instance.
3. Repeat the above iteration 1000 times to calculate the mean and standard error of the performance.

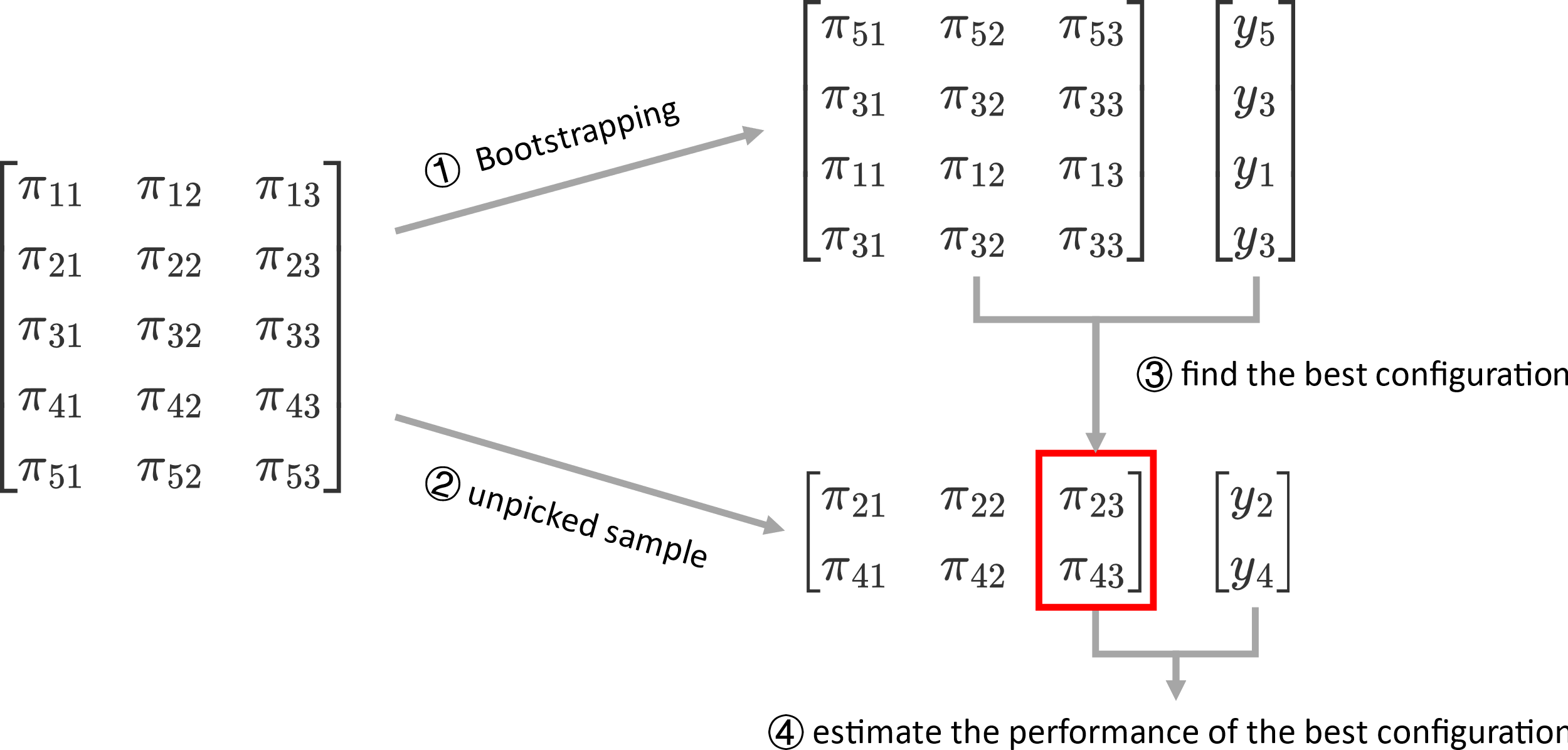

Figure S3. Example of one iteration of BBC-CV. (1) the five out-of-sample prediction from three configuration are split into two groups; (2) Pick the 1nd, 3rd and 5th data instances to estimate the performance of each configuration. Since this is a bootstrap step and allow replacement, the 3rd data instance can be picked twice. The 3rd configuration performs the best in this example; (3) Finally, use the remaining 2st and 4th data instances to estimate the unbiased performance.
